# A Crispr Knockout Screen Identifies Foxf1 as a Suppressor of Colorectal Cancer Metastasis that Acts through Reduced mTOR Signalling

**DOI:** 10.1101/613356

**Authors:** Lennard Lee, Connor Woolley, Thomas Starkey, Luke Freeman-Mills, Andrew Bassett, Fanny Franchini, Lai Mun Wang, Annabelle Lewis, Roland Arnold, Ian Tomlinson

**Affiliations:** Institute of Cancer and Genomic Sciences, College of Medical and Dental Sciences, University of Birmingham, Edgbaston, Birmingham B15 2TT, UK; Wellcome Trust Sanger Institute, Cambridge CB10 1SA; Kennedy Institute of Rheumatology, Oxford OX3 7FY; Department of histopathology, John Radcliffe Hospital, OX3 9DU; Cancer Gene Regulation Laboratory, Oxford Centre for Cancer Gene Research, Wellcome Trust Centre for Human Genetics, University of Oxford, Roosevelt Drive, Oxford OX3 7BN, UK

## Abstract

**Introduction:** A greater understanding of molecular mechanisms underlying metastasis is necessary for development of new strategies to prevent and treat cancer.

**Methods:** We performed a genome-wide CRISPR/Cas9 knockout screen in MC38 colorectal cancer (CRC) cells transplanted orthotopically into mice to identify genes that promote metastasis. We undertook focussed molecular analyses to identify mechanisms underlying metastasis.

**Results:** The screen identified several gene knockouts over-represented in lung metastases, including Dptor (mTOR signalling) and Foxf1 (gastrointestinal tumour predisposition). We validate that loss of Foxf1 promotes metastasis, increased Foxf1 expression restrained cellular migration in-vitro and human CRC metastases express lower Foxf1 than paired primary tumours. Analysis of gene expression changes downstream of Foxf1 identified increased mTOR signalling as a possible mechanism of metastasis caused by Foxf1 loss, consistent with Dptor identification. We confirmed this mechanism demonstrating that mTOR inhibitor sirolimus reduced lung metastasis burden in xenografts.

**Conclusion:** Mesenchymal Foxf1 plays a major role in intestinal development. We have shown for the first time, through an unbiased genetic screen, that reduced epithelial Foxf1 results in raised mTOR signalling and metastasis.

**Authorship statement:** Lennard Lee-study concept and design, acquisition of data, analysis, interpretation of data, drafting of the manuscript, statistical analysis and obtained funding. Connor Woolley-acquisition of data, analysis and interpretation of data. Thomas Starkey-acquisition of data, analysis, interpretation of data, drafting of the manuscript. Luke Freeman-Mills-interpretation of data. Andrew Bassett-technical and material support. Fanny Fanchini-technical support. Lai Mun Wang-acquisition of data and study supervision. Annabelle Lewis-study supervision. Roland Arnold-analysis, interpretation of data, statistical analysis. Ian Tomlinson-study supervision and critical revision of the manuscript.

**Conflict of Interest:** The authors whose names are listed above declare that they have no conflict of interest.

## Introduction

Metastatic disease is the principal cause of death in patients with colorectal cancer (CRC) and more than half of CRC patients present with metastases (1). In order to metastasise, a cancer cell requires changes in cellular function, including the development of cell-autonomous traits like migration and invasion, but also the ability to modulate interactions with host mesenchymal, endothelial and immune cells. Metastatic cells must also survive in blood, lymph or other media such as peritoneal fluid. There is a partial molecular understanding of the processes that govern CRC metastasis, with metastasis-promoting identified from mouse models for *BRG1* (2), *MACC1* (3), *HIF1A* (4), *UBE2V1* (5), *CPT1A* (6) and *EPCAM* (7).

Genomic studies searching for metastasis-specific driver mutations have had limited success in humans (8,9). Unravelling the complexity of metastasis may therefore require alternative strategies that complement human cohort-based approaches. Genetic screens in CRC animal models have afforded a large-scale, unbiased and comprehensive approach to identifying processes that promote or retard the early stages of colorectal tumorigenesis (10) (11) (12). The strengths of such an approach principally lie in avoiding the genetic noise observed in human cancers, which may result from host factors (age, lifestyle, socio-economic), tumour factors (location, size, degree of invasion) and treatment factors.

To date, there have been no genetic screens in CRC mouse models to identify determinants of metastasis. In this study, we report the results of the first genome-wide agnostic screen for gene knockouts that promote CRC metastasis in a novel orthotopic mouse model based on colonoscopic delivery of CRC cells. Our findings highlight a role in metastasis for the *Foxf1* gene and mTOR pathway, results that are validated in human CRCs, and open new avenues for research into CRC metastasis.

## Methods

### CRISPR GeCKO genetic screen

Generation of the GeCKOv2 mouse lentiviral library A (GeCKO-A) was performed as previously described (13,14). Model-based Analysis of Genome-wide CRISPR/Cas9 Knockout program (MAGeCK) (15) was utilised identify gRNAs and genes targeted. gRNAs not detected were given a score of 0. Remaining gRNAs for each mouse were divided into quintiles based on read count, with 1 the lowest quintile. A final score was computed for each gene by summing up all the quintile numbers for all guides corresponding to that gene for all mice.

### Transcriptomic analyses

RNA sequencing libraries were prepared using the QuantSeq 3’ mRNA library prep kits *(Lexogen)* and sequencing performed on an Illumina Hiseq 4000 with 75bp paired end reads.

### Generation of knockout and CRISPR/Cas9 Synergistic Activation mediators

Knockout of *Foxf1* in MC38 was achieved through the use of three gRNAs that were cloned into pLentiCRISPRv2 vector. CRISPR/Cas9 Synergistic Activation mediators (CRISPR/Sam) were cloned into the lenti sgRNA(MS2)_puro backbone vector and transduced into stable cell lines expressing dCAS-VP64 and MS2-P65-HSF1.

### Public repositories of human CRC cohort data

Data were obtained from the Gene Expression Omnibus, using the datasets GSE41258 (16), GSE6988 (17) and GSE22834 (18). Expression data from The Cancer Genome Atlas (TCGA) colorectal cancer cohort, was obtained directly from the Broad Institute website (https://gdac.broadinstitute.org).

## Results

### CRISPR/Cas9 knockout screen to identify metastasis suppressor genes

A whole-genome CRISPR/Cas9 knockout screen (GeCKO-A) was utilised to identify genes that promoted CRC metastasis when knocked out. The mouse CRC cell line MC38 was chosen as it forms rapidly growing tumours when transplanted into the colon of mice with low rates of metastasis. Cells stably expressing firefly luciferase (MC38-Zsgreen-Luc2) were used to facilitate bioluminescence-based *in vivo* imaging.

MC38-Zsgreen-Luc2 cells were transduced with the GeCKO-A library, consisting of 67,405 guide RNAs (gRNAs) targeting 20,611 protein-coding genes. Using a precision endoscopic technique, transduced cells were transplanted in bulk submucosally in the descending colon of Nod/Scid gamma (NSG) mice (Figure 1). 6 test mice were generated in two independent, temporally separated transduction replicates. Untransduced MC38 cells were used to produce four control mice in similar fashion.

**Figure 1.**
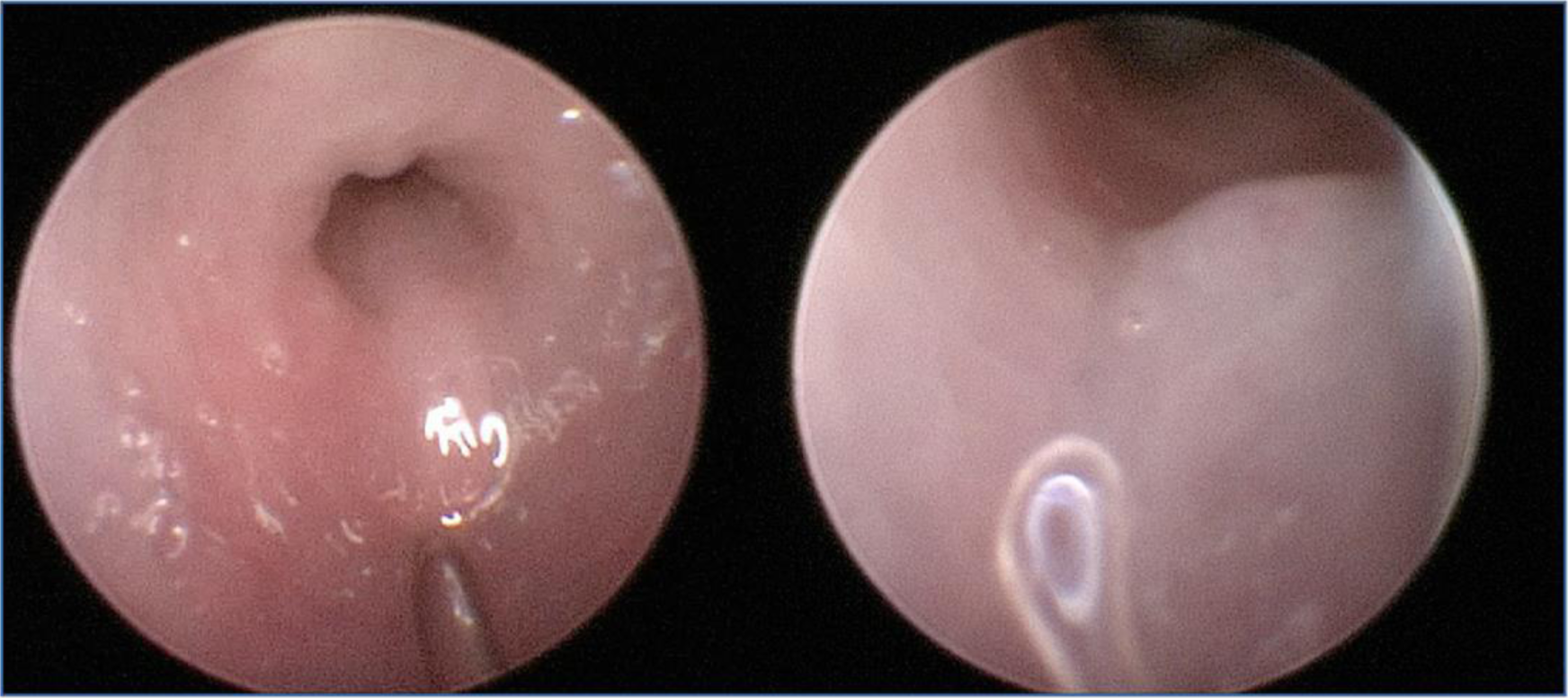
Metastases derived from orthotopic injection of MC38 cells from the CRISPR knockout screen. Endoscopic image demonstrating technique for orthotopic microscopic transplantation of MC38 cells. The photographs show the lumen of the descending colon of mice, with the left-hand image demonstrating submucosal injection and the right-hand image showing the resulting growth of a primary tumour 7 days later.

Transplantation of MC38 cells induced colonic tumours in all mice, causing intestinal obstruction from the third week after transplantation. Metastases were detected in the lungs of all mice, with increased burden in test animals than controls (independent sample t-test, p=0.049). Liver metastases were infrequently observed, (3 test mice, 1 control). At the assay resolution, no metastases were found in bones, brain, spinal cord, spleen or kidneys.

### Identification of gene knockouts associated with lung metastasis

We searched for gene knockouts that promoted metastasis, cognisant of the fact that their effects may have involved any stage(s) of tumorigenesis *in vivo*, from primary tumour, through dissemination, to growth of the metastasis in the lung. To identify gRNAs associated with metastasis, the lungs were resected whole from each mouse, DNA extracted and targeted next-generation sequencing performed. Pre-transplantation GeCKO-A-transduced MC38-Zsgreen-Luc2 cells were used as baseline. The baseline cell population had 96.6% representation of the GeCKO-A library (63,686/65,960 gRNAs). Lung metastases showed reduced library representation (average 13.7%) likely resulting from a combination of founder effects, drift and negative selection.

In lung metastases, gRNAs for a median of 2,549 genes were identified per test animal. As observed in a previous publication using a CRISPR screen (19), read counts for gRNAs were highly skewed, with some guides dominating the sequencing and collecting thousands of reads, while others had very few reads. The significance of high read counts is unclear, and may result not only from selection, but also from PCR amplification during sequencing or early exponential outgrowth of metastases. We reasoned that, despite a potentially large random component to metastasis (for example, cells that hitch-hike with others during bulk migration), gRNAs targeting genes that promote metastasis when inactivated would fulfil the following criteria: (i) an enrichment in sequencing reads from lung lesions; (ii) reproducibility between different gRNAs targeting the same gene; and (iii) reproducibility between mice.

Overall, there was strong evidence for concordance of gRNA enrichment between mice (Supp. Table 1). For each mouse, guides were ranked by normalised read count and the resulting distribution was divided into quintiles (0-4). Finally, a Metastasis Screen Score (MSS) was computed for each gene by summing the quintile numbers for all guides for that gene across all test mice.

**Table 1.**
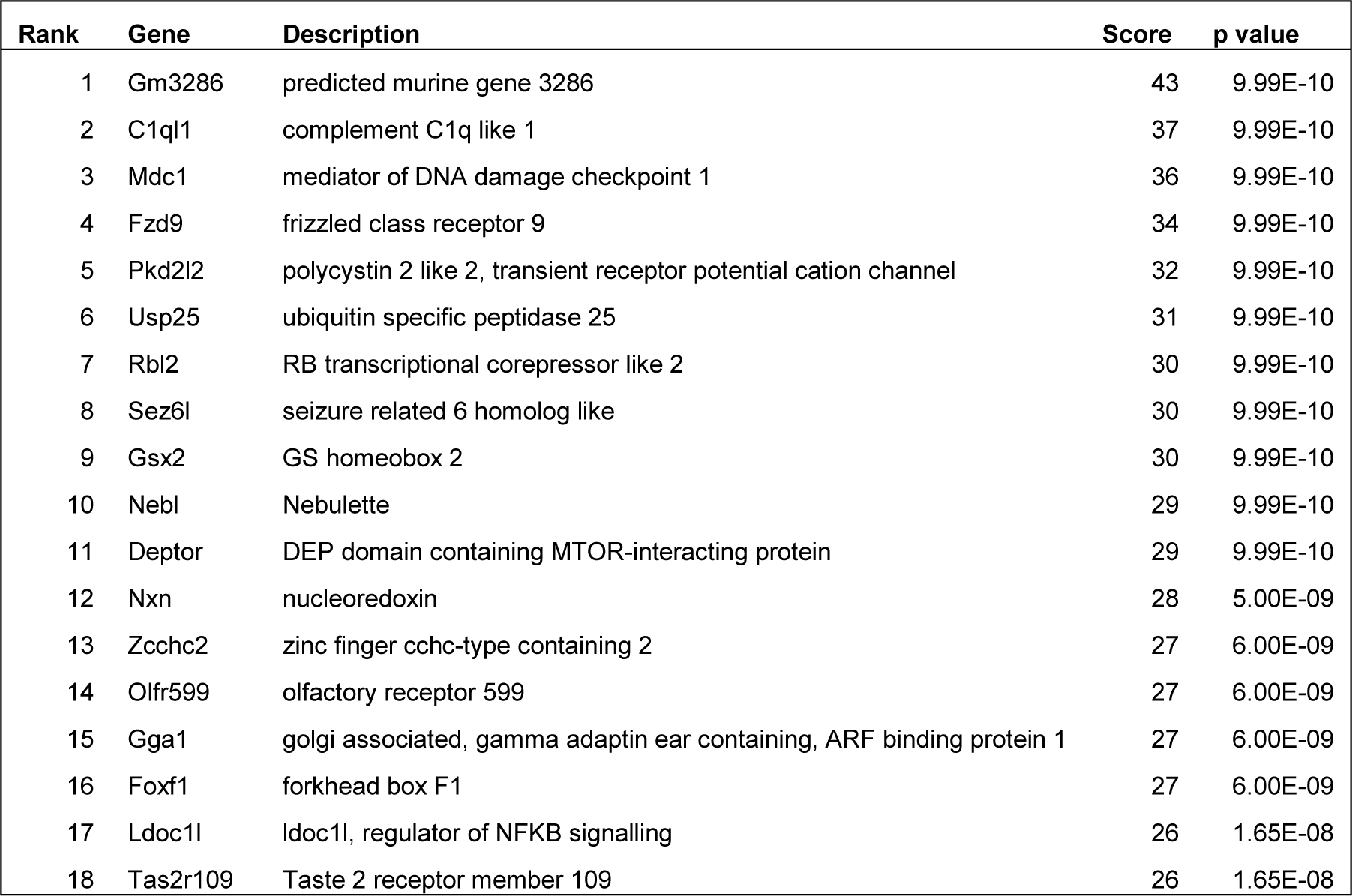
Genes enriched in CRISPR/Cas9 from metastasis screen. Genes were selected with a final score of greater than 25 and represent gRNA that were over-represented in the metastatic lung lesions. p values represent the expectation value of the observation based on randomisations on the dataset.

We prioritised for further analysis genes with an MSS >25, resulting in a list of 18 genes (Table 1, Supp. Table 2). These genes are respectively involved in DNA damage response (*Mdc1),* Wnt signalling *(Fzd9, Nxn),* histone modification *(Rbl2),* mTOR signalling *(Dptor)*, and gut development and gastrointestinal tumour predisposition (*Foxf1*). Of this priority gene list, we chose *Foxf1* for further analysis owing to its potentially specific role in CRC biology, as polymorphisms at this locus are involved in predisposition to gastric (20), oesophageal (21) (22) and colorectal cancers (23).

### Foxf1 restrains metastasis in vivo

We validated findings of the CRISPR/Cas9 screen using three new gRNAs targeting *Foxf1*. The resulting MC38-Foxf1-ko cells exhibited multiple *Foxf1* coding insertion/deletion mutations, with a concurrent average decrease in *Foxf1* mRNA levels of >75% (Supp. Figure 1). Following orthotopic transplantation into NSG mice, all developed colonic tumours (n=8). There was no significant difference in primary tumour size between knockdown and wild type cells, but lung metastasis burden was increased 2.96-fold in the former (p=0.01) (Figure 2). These findings not only validate results of our screen, but also demonstrate that *Foxf1’s* effect on CRC metastasis does not principally result from limiting proliferation of the primary tumour.

**Figure 2.**
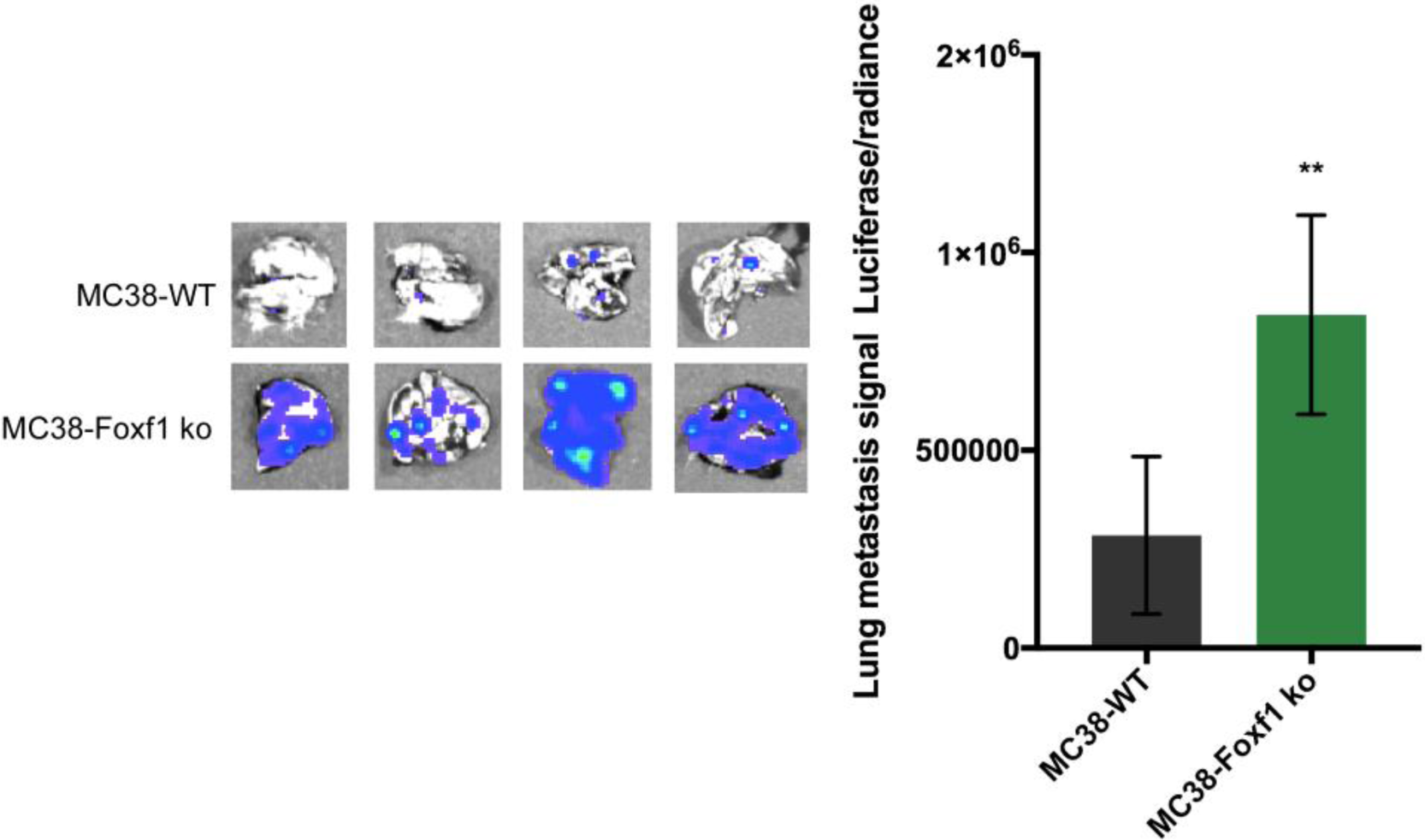
Knock out of Foxf1 in MC38 using a CRISPR/Cas9 strategy causes an increased burden of lung metastasis after tail vein injection. Images show resected whole lung specimens and *ex vivo* bioluminescent imaging. Wild type MC38 demonstrates the typical presence of small, isolated lesions. Foxf1 knockdown using three new validation sgRNAs, resulted in an increase in number and size of lesions, with a 2.96-fold increase in lung metastasis burden based on luciferase assays (p=0.01).

### FOXF1 re-expression decreases cellular migration

Although *Foxf1* mRNA levels are relatively high in MC38, FOXF1 expression is frequently reduced or lost in human CRC cell lines (24) (Supp. Figure 2). Thus, rather than knocking out *FOXF1* in these cells, we re-expressed it using CRISPR/Cas9 Synergistic Activation Mediator (CRISPR/Sam). In the four CRC lines (HT29, HCT116, LS174T, SW480), the resulting *FOXF1* mRNA levels were comparable to normal colonic tissue (Supp. Figure 2). The effect of re-expressed *FOXF1* on cellular migration was determined using a Boyden migration assay. This demonstrated a decrease in migration in all four lines tested, with a mean decrease of migratory cells of 59% (two sided t-test, p<0.01 for all; Supp. Figure 3). These results are consistent with alterations in cellular migration being a contributory factor to the restraining effect of *FOXF1* on metastasis.

**Figure 3.**
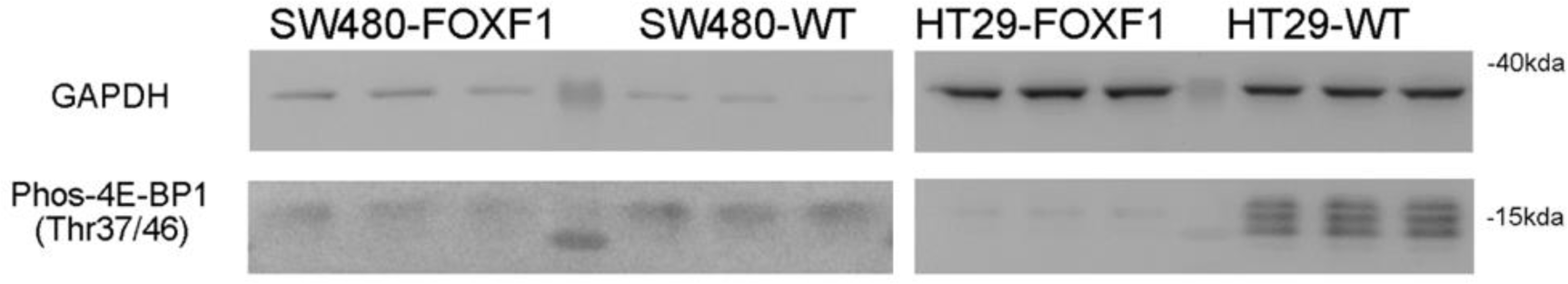
Western blot demonstrating relationship between FOXF1 and mTOR activity. Representative western blot demonstrating inhibition of mTOR effector protein phospho-4E-BP1 following *FOXF1* re-expression in CRC cell lines SW480 and HT29. There was consistent down-regulation of phosphorylated 4E-BP1 (60% and 96% respectively, p<0.005).

### FOXF1 expression levels are inversely correlated with mTOR signalling

To gain biological insights into mechanisms associated with changes in *FOXF1* expression in CRCs, we performed an analysis of publicly available data from The Cancer Genome Atlas (TCGA), comprising of transcriptomic profiles of 255 CRCs. Tumours were separated into those expressing high *FOXF1* and low *FOXF1* by bifurcation of RSEM count data at the mean (25). We identified 2,853 highly differentially expressed genes (FDR q<5×10^-5^) between *FOXF1*-high and *FOXF1-*low CRCs (Supp. Table 3). The overlap with MSigDB hallmark gene sets was assessed, and identified that *FOXF1*-high expressing CRCs had down-regulation of “Myc Targets” (p=9.65×10^-74^), “Oxidative phosphorylation (p=2.70×10^-65^), “E2F targets” (p=4.76×10^-49^) and “mTORC1 signalling” (p=7.51×10^-31^, Supp. Figure 4), and up-regulation of “Epithelial-Mesenchymal Transition (EMT)”, (p=1.21×10^-68^) (Supp. Table 4). Identification of negative association between *FOXF1* and *RB/E2F* signalling (26) and positive association with epithelial-mesenchymal transition confirmed previous reports (27), but the potential relationship between high *FOXF1* and inhibition of mTOR signalling is novel and has not been shown previously.

**Figure 4.**
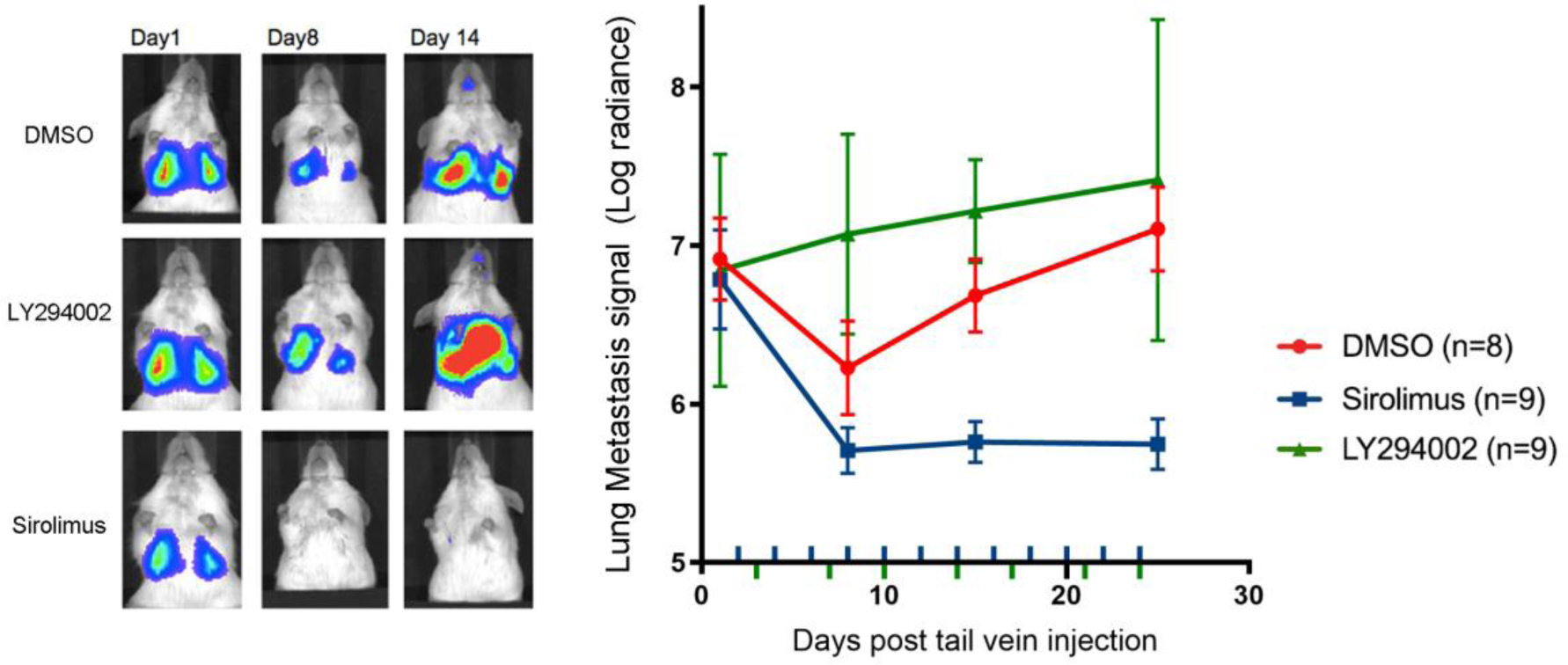
Effects of mTOR and PI3K inhibition on lung metastasis outgrowth in a mouse tail vein metastasis assay. A. Representative *in vivo* bioluminescence images of lung metastases following tail vein injection of human CRC cell line LS174T in a single mouse given either DMSO, LY294002 (PI3K inhibitor) or Sirolimus (mTOR inhibitor). At day 1, LS174T cells lodge in the lung parenchyma, at day 8 there is a mild decrease signal due to a proportion of cells not having the capability to establish in the lung parenchyma, from day 15 onwards, there is metastatic outgrowth. B. Graph demonstrating the lung metastasis burden in mice given DMSO (n=8), LY294002 (n=9) or Sirolimus (n=9). Lung metastasis burden was determined by *in vivo* bioluminescence imaging. Error bars denote standard deviation. Coloured ticks represent dosing intervals. In the DMSO control, at day 1, LS174T cells lodge in the lung parenchyma, at day 8 there is a mild decrease signal due to a proportion of cells not having the capability to establish in the lung parenchyma, from day 15 onwards, there is metastatic outgrowth. In mice that received Sirolimus, there was a significant decrease in the lung metastasis burden at day 7, 15 and 21 compared to those that received DMSO (independent sample t test, p=0.007). In mice that received LY294002, there were no significant differences in lung metastasis burden at day 7, 15 and 21 compared to those that received DMSO.

To identify gene sets that might be directly influenced by FOXF1 expression, we interrogated the transcription factor database, TRANSFAC, to search for predicted transcription factor targets for FOXF1 using a position weight matrix algorithm. This identified 1,508 potential FOXF1 target genes with predicted FOXF1 binding sites (28). MsigDB hallmark gene set analysis showed significant overlap with “epithelial-mesenchymal transition” (FDR q-value=6×10^-5^) and “mTORC1 signalling” (FDR q value=2.3×10^-5^) (Supp. Table 5A, 5B).

We then re-expressed *FOXF1* in 7 CRC cell lines (HT29, HCT116, LS174T, SW480, T84, GP2D, SW1222) using CRISPR-Sam and transcriptomic profiles were analysed. mRNA-based gene set enrichment analysis (GSEA) identified 15 Hallmark signalling networks that were impacted following re-expression of *FOXF1* (Supp. Table 6), one of which was mTORC1 signalling, which was diminished, validating our bioinformatic analyses of the TCGA and TRANSFAC data. This observation was consistent with our initial finding that *Deptor*, an mTORC inhibitor, was one of the top-ranked hits in our CRISPR/Cas9 screen.

We sought to validate this observation at a protein level. We therefore utilised FOXF1 re-expressing SW480 and HT29 cell lines. In both lines, FOXF1 re-expression caused consistent down-regulation of phosphorylated 4E-BP1, an effector of mTOR signalling (60% reduction and 96% reduction respectively, p<0.005 for both; Figure 3). We also analysed protein levels of the mTORC1 scaffolding protein, RPTOR (Raptor), as an additional marker of mTOR activity. Western blotting demonstrated that Raptor expression was reduced by an average 60% in FOXF1 re-expressing cells (p<0.05; Supp. Figure 5). These results are consistent with a function of FOXF1 being transcriptional repressor of mTOR activity.

**Figure 5.**
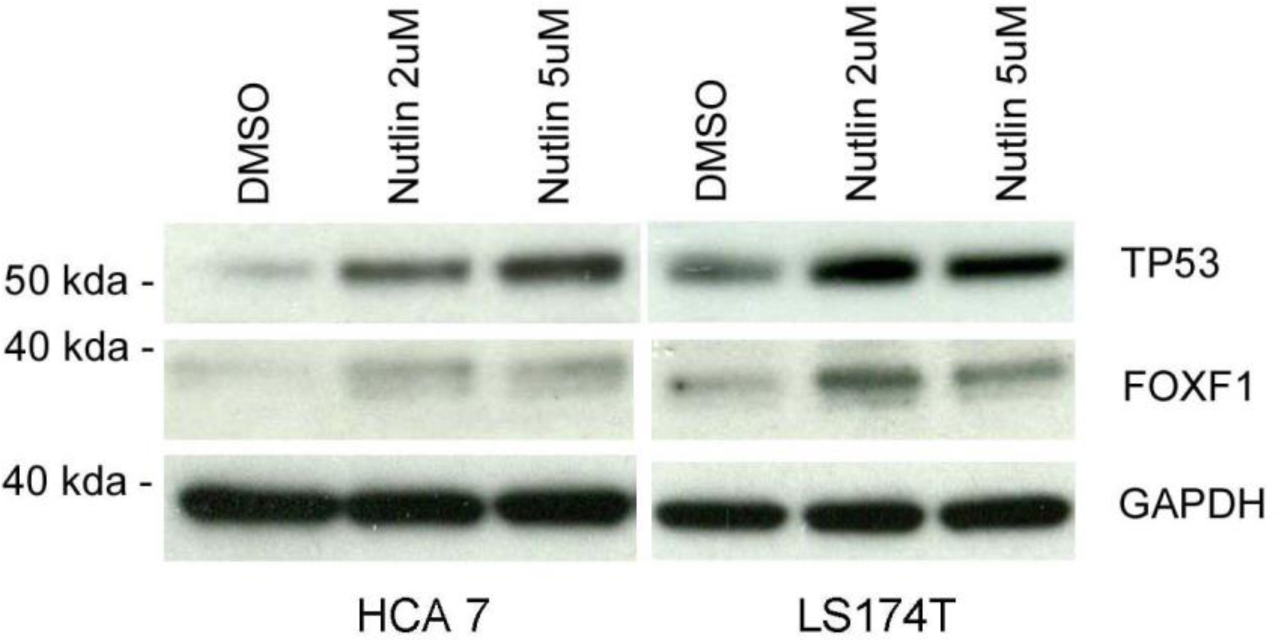
Representative western blots demonstrating that Nutlin-3-mediated increased expression of TP53 results in higher FOXF1 expression. The assay shows an average 2.4-fold increase in FOXF1 expression (p=0.002) in HCA7 (*TP53* mutant/wildtype heterozygote) and LS174T (*TP53* wildtype).

In most cancers, especially CRC, mTOR signalling is constitutively active, and expression of its inhibitor *Deptor* is low. In agreement with our CRISPR/Cas9 screen data (see above), low *Deptor* expression has been reported to correlate with nodal metastasis in CRC patients (29). We also noted that MC38 cells have ∼4-fold higher *Deptor* expression and a lower metastatic potential than human CRC lines (data not shown). We therefore suppressed *Foxf1* expression in MC38 using a mini-CRISPR/Cas9 gRNA pool. In comparison with parental cells, we identified 119 differentially expressed genes (FDR q <5×10^-5^). Gene set enrichment analysis identified a significant association between *Foxf1* knockout and increased mTORC1 signalling and Epithelial Mesenchymal transition (both FDR q =0.01; Supp. Table 7).

### mTOR inhibition reduces the ability of circulating cancer cells to seed and grow

The effect of mTOR inhibition specifically at the level of CRC metastatic lesions has not been fully described. Furthermore, the dependence of metastatic mTOR activity on upstream PI3K signalling is unclear.

In order to analyse CRC metastases and exclude effects occurring in the primary cancer (metastasis initiating effects), we injected human LS174T CRC cells into the lateral tail veins of NSG mice, which then seed in the lungs, and assessed subsequent metastatic development. Following metastatic implantation, mice then received the mTOR inhibitor Sirolimus (n=9), the PI3K inhibitor LY294002 (n=8), or vehicle (n=8). Sirolimus significantly inhibited outgrowth of metastases (94% lower at day 25; t-test, p=0.007), whereas no significant differences were observed with LY294002 (Figure 4). These results confirm the importance of mTOR signalling in enabling distant metastasis outgrowth, whilst showing independence of PI3K signalling.

### FOXF1 expression is controlled by TP53 response elements

*TP53* is a tumour suppressor and there is evidence from breast cancer experiments that its inactivation can contribute to cancer metastasis (30) (31), TP53 activation has an inhibitory effect on mTOR activity in MEF cells (32) and *FOXF1* expression has been shown to be induced in a TP53-dependent manner through a TP53 response element in *FOXF1* intron 1 (33).

We sought to validate these findings, and demonstrate that *TP53* inhibits mTORC activity at least in part through *FOXF1*. We activated TP53 using Nutlin-3, an MDM2 inhibitor, in two human CRC cell lines (HCA7, LS174T), which have at least one intact *TP53* allele and low/normal *TP53* expression. Western blot analysis demonstrated that Nutlin-3 led to increases in *TP53* expression, with a corresponding 2.4-fold increase in *FOXF1* expression (p=0.002; Figure 5).

This experiment shows positive correlation between *TP53* expression and *FOXF1*. This suggests that TP53 inactivation results in lower FOXF1 levels and we propose that this could then increase metastasis through increased *mTOR* signalling. This model is consistent with the observation in our original CRISPR knockout screen that *Trp53* knockout promotes metastasis (Supp. Table 2).

### Low FOXF1 expression is observed in human CRC metastases

Having established the importance of FOXF1 in mediating mTOR inhibition and retarding metastasis in animal models, *in vitro* experiments and publically available datasets, we analysed *FOXF1* mRNA expression in human CRC metastases. If *FOXF1* does restrain metastatic outgrowth, we might expect decreased *FOXF1* expression in metastatic lesions compared to the primary tumours. Conversely if this was not observed, this could suggest that *FOXF1* loss acts as a metastatic promoter principally at the primary site. The publically available dataset GSE41258 (16) comprising of 186 primary CRCs and 67 distant metastases (47 liver and 20 lung) was investigated. *FOXF1* was significantly decreased in lung metastases (1.72-fold, p<0.0011) and liver metastases (2.2-fold, p<0.001) compared to the primary tumours (Supp. Figure 6A). This was validated in the GSE41258 dataset consisting of CRC liver metastases (Supp. Figure 6B).

**Figure 6.**
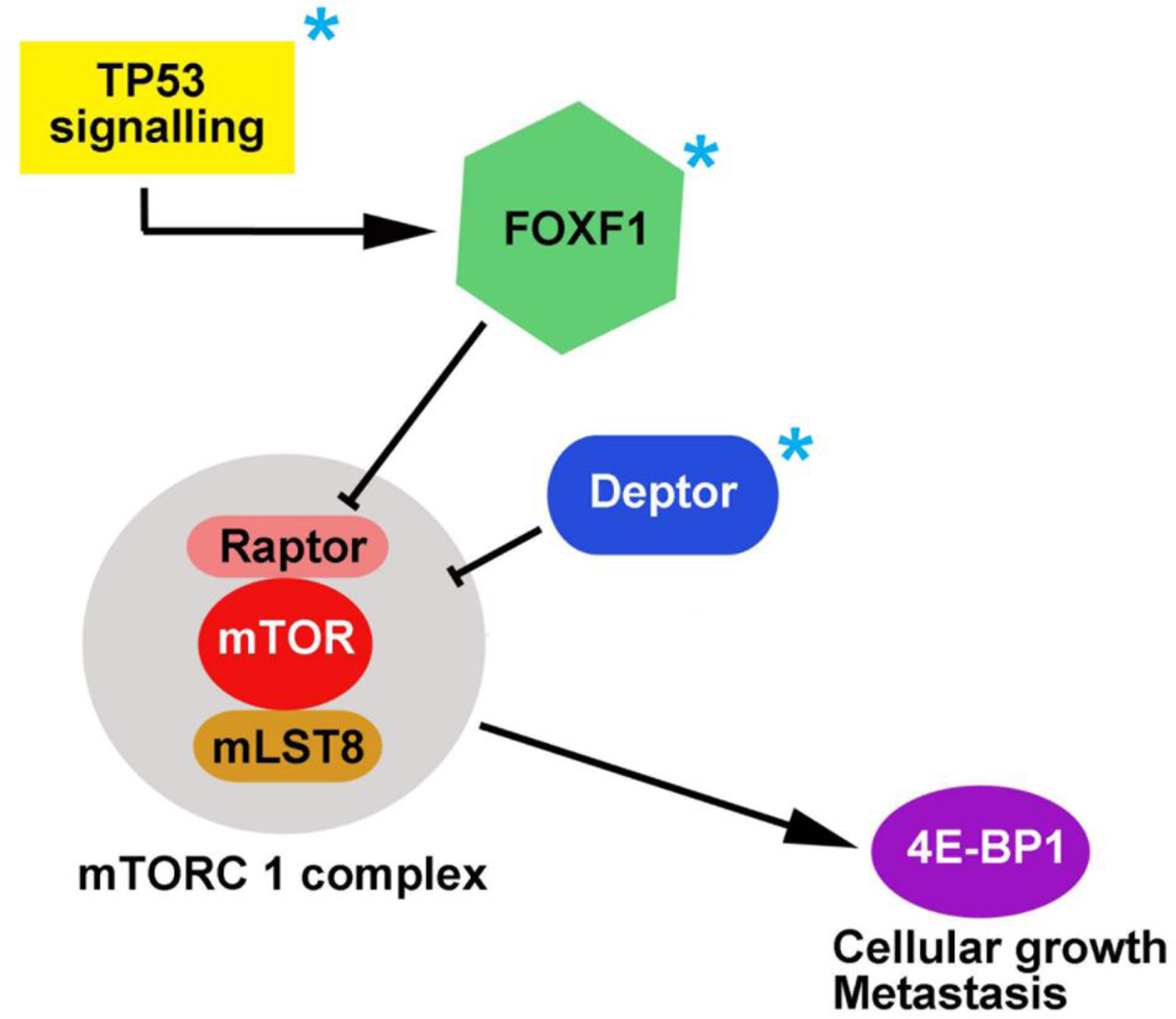
Hypothesised mechanisms linking FOXF1 loss of function to metastasis. Blue stars indicate genes which were enriched from CRISPR/Cas9 knockout screen.

Having established a link with decreased *FOXF1* expression in distant metastases (liver and lung), we proceeded to analyse whether this is also observed in local lymph mode metastases. We analysed our own data from the *METALLIC* sample set *(Metastatic lymph nodes in colorectal cancer)*. This sample set consists of paired stage 3 primary tumour and lymph node metastases for 41 patients. We found that expression of *FOXF1* mRNA was significantly lower in lymph node metastases than their paired primary CRCs (paired t-test, p=0.038; Supp. Figure 6C).

In combination, these observations suggest that relative to primary tumours, reduced *FOXF1* is a common feature of CRC metastases.

## Discussion

There has been growing awareness of the ability of cancer cells to re-activate embryonic developmental processes. Forkhead Box (FOX) family proteins are evolutionarily conserved transcription factors (TFs), crucial in embryonic development that regulate a multitude of genes involved in the cell cycle and differentiation. There is increasing evidence that FOX proteins are dysregulated in cancer, contributing to tumour initiation, progression and metastasis. (34–36).

In this contribution, we have identified several genes with evidence of modulating metastasis risk including genes involved in the DNA damage response (*Mdc1*), Wnt signalling (*Fzd9* and *Nxn*), histone modification *(Rbl2),* mTOR signalling *(Dptor)*, and gastrointestinal tumour predisposition (*Foxf1*). Our experimental approach has several strengths, being genome-wide, with direct assessment of metastatic burden and the generation of sub-mucosal tumours within the descending colon through our novel endoscopic transplantation approach.

We focussed on *FOXF1* owing to recent evidence suggesting a role in human CRC susceptibility (23). *FOXF1* is normally a mesenchyme-specific TF, and a likely target of hedgehog signalling and is reported to have a role in the tumourigenesis of breast carcinoma (26), osteosarcoma (37) and rhabdomyosarcoma (38). *FOXF1* act as a transcriptional repressor, suppressing cell cycle regulatory proteins (38), although a role as an activator in certain situations remains possible.

We confirmed that loss of *FOXF1* promotes metastasis in focussed assays. We used multiple methods, different cell lines and gRNA from the original screen, including re-expression and knock down of *FOXF1* and whole transcriptomic analyses, to determine potential mechanisms of *FOXF1’s* action on metastasis and ensure our results were robust. Of the large number of gene signalling pathways perturbed by *FOXF1*, we show for the first time that down-regulation of the mTOR pathway was most apparent. This is consistent with observations from the CRISPR/Cas9 screen that loss of function of the negative mTOR regulator *Deptor* also promotes metastasis. The negative association between *FOXF1* expression and mTOR activity in human CRCs and down-regulation of *FOXF1* in metastatic human CRCs provides further support for this new mechanism.

*TP53* mutation is associated with metastasis and has been linked to *FOXF1* expression (33). We were able to confirm this and hypothesise that *FOXF1* forms a bridge linking *TP53* and mTOR signalling. Mutated or epigenetically silenced *TP53* would lead to decreased *FOXF1*, which in turn would lead to increased mTOR signalling and promote metastasis (Figure 6).

There are some caveats that must be borne in mind. Firstly, selection of “top” genes in such screens is inevitably heuristic. Secondly, due to the inherent nature of CRISPR genetic screens, they are prone to false positive or negative results dependent on gRNAs performance and they do not identify genes/networks controlled by post-translational modification. Finally, metastasis may well be a process with a large stochastic element and our data suggest that multiple sub-clones from the primary cancer may colonise and grow in the lung.

In summary, this CRISPR screen approach has enabled us to identify genes/pathways likely to be of importance in the spread of CRCs. By identifying *FOXF1,* the mTOR pathway and several other candidate genes that modulate the likelihood of metastasis, we have developed a greater understanding of metastasis biology with the potential to develop new therapeutic strategies.

## Acknowledgements

LL was supported by an MRC Clinical Research Fellowship and NIHR BRC Clinical Career Development fellowship and received project grants from the Guts UK Charity (previously known as Core UK) and Cancer Research UK. CW and much of the experimental work was supported by ERC Senior Award EVOCAN to IT. AL was supported by MRC Grant. We acknowledge the Wellcome Trust Core Award to the Wellcome Trust Centre for Human Genetics (090532/Z/09/Z).

